# Discovery of Thermodynamic Control Variables that Independently Regulate Healthspan and Maximum Lifespan

**DOI:** 10.1101/2024.12.01.626230

**Authors:** Kirill A. Denisov, Jan Gruber, Peter O. Fedichev

## Abstract

The question, “Can aging be modified, delayed, or reversed?” has profound social and economic implications for rapidly aging societies today. Interventions, ideally, would intercept functional decline and extend healthspan by delaying late-life morbidity (known as “squaring the curve”). These have proven elusive, but examples of differential aging in the animal world abound, suggesting aging itself is a malleable process. We present a novel multi-scale theoretical framework for entropic aging, and apply it to recently published DNA methylation data from 348 evolutionarily distant mammalian species. Our analysis identified modules or correlated DNA methylation changes associated with reversible pathway activation in key biological processes. We discovered a single species-dependent scaling factor controlling the magnitude of fluctuations across biological pathways. It acts as the organism’s “effective temperature”, quantifying intrinsic biological noise within networks and is unrelated to physical body temperature. Furthermore, we find a distinct stochastic damage signature and an associated extreme value (Gumbel) distribution of activation barriers controlling site-specific damage rates of individual CpG sites. This implies that aging is driven by rare, high-energy transitions on rugged energy landscape, most likely simultaneous and hence practically irreversible failures in highly redundant systems. While the overall rate of damage accumulation and hence the maximum lifespan does not depend on the effective temperature driving the noise in leading pathways, effective temperature does influence both initial mortality rate and the mortality rate doubling time – thereby shaping the survival curve. Lowering effective temperature must, therefore, be a promising Geroscience strategy, aimed directly at squaring the curve of aging. The example shows that targeting the thermodynamic forces driving mammalian aging may provide powerful strategies for the development of truly meaningful interventions to combat aging in humans.

## I. INTRODUCTION

It has been said that physics is the law, while everything else is merely a recommendation [1]. Although evolution functions as a powerful optimization algorithm, it too has to operate within the limits set by the laws of physics. These constraints give rise to structures and properties that emerge consistently across large sections of the tree of life. Understanding these universalities can therefore provide insights that differ fundamentally from those gained through microscopic investigations of the same phenomenon. Allometric scaling laws are perhaps the earliest demonstration of this principle, exemplifying how physical constraints – specifically, the minimization of dissipation driven by natural selection – prevent straightforward isometric scaling of metabolism with body size. The consequence is an exponent of 3/4 in the scaling relationship between body mass and metabolism, a relationship known as Kleiber-West law [2, 3]. Applying this relationship to the first law of thermodynamics (energy conservation) provides an elegant model for ontogeny—the development and growth of an organism’s body [4].

In G. West’s model of growth, there is no aging— the fully developed organism represents a steady state solution. However, in [5], we observed that the stability of an organism, as measured by its recovery rate (or resilience) declines linearly with age in humans. Linear age-dependent decline in physiological resilience is a general feature of aging that is typically present significantly before overt disease or frailty can be detected [6]. Declining physiological resilience is therefore naturally explained as consequence of linear damage accumulation, that is, the linear decline in homeodynamic performance results from a large number of independent, stochastic state transitions or macromolecular modifications, each having small, typically detrimental effects on the performance of random nodes in these networks. In more abstract language, the linear decline in homeodynamic resilience is the consequence of an age-dependent, linear increase in configurational entropy [7, 8]. Therefore, aging may be most accurately understood within the framework of the second law of thermodynamics, particularly in relation to entropy increase and macroscopic reversibility in systems with limited microscopic control.

In this study, we introduce a novel multi-scale theoretical framework to understand entropic aging at both the whole-organism (macroscopic) and pathway activation (microscopic) levels and apply it to newly published DNA methylation (DNAm) data from 348 mammalian species spanning large phylogenetic distances [9]. Using linear matrix factorization, we identify modules of correlated DNAm features separable into two qualitatively distinct classes: one displays the hallmark signatures of stochastic damage accumulation, while the other comprises signals of reversible pathway activation that map to key biological processes (organism-wide pathways). Focusing on the former, we derive an empirical damage accumulation rate (DAR) for each species and demonstrate that this quantity is inversely proportional to both the development time and renewal rate (*r*) from G. West’s theory of ontogeny. This finding links microscopic damage accumulation dynamics to macroscopic scaling constraints, strongly supporting the idea that aging is tightly coupled to biological renewal dynamics.

We also observed that the biological noise, quantified by the magnitude of fluctuations in activations of leading biological pathways is controlled by a single factor, best interpreted as the “effective temperature”, *T*_eff_. The effective temperature is a concept borrowed from nonequilibrium thermodynamics and relates the magnitude of fluctuations with the power of dissipation through the fluctuation-dissipation theorem [10]; *T*_eff_ functions as a universal scale for the noise in mammalian regulatory networks; it is different from and should not be confused with the physical body temperature.

At the microscopic level, we propose a model that links the kinetics of stochastic damage accumulation with the structural properties of the regulatory network – specifically the distribution of the activation barriers separating microscopic configurations of networks. We mapped the empirical exponential distribution of the site-specific rates of change in DNAm to our model and found that the energetic barriers protecting individual methylation sites follow a Gumbel distribution, suggesting a form of universality in the aging process rooted in extreme value statistics. This observation suggests that these rare, high-impact events play a crucial role in the aging process. Consequently, aging emerges as a thermodynamically irreversible phenomenon, as reversing these rare stochastic events would demand an impractical degree of control at the microscopic level.

Our analysis suggests that critical actuarial aging parameters—including the initial mortality rate and the Gompertz exponent—are highly sensitive to the effective temperature, thereby determining the difference between maximum and average lifespan. Furthermore, targeting the effective temperature must represent a powerful strategy to extend healthspan by “squaring the survival curve.” Consequently, we advocate for the development of a new class of longevity therapeutics that target these thermodynamic forces underlying mammalian aging at the macroscopic level. We assert that this approach is the only viable strategy for enabling truly meaningful interventions to combat aging in humans.

## II.RESULTS

### A. Background on allometric scaling, energy conservation and the theory of ontogenic growth

The basic metabolic rate *B* scales with body mass *m* according to the Kleiber-West allometric scaling law: *B*(*m*) = *αm*^3*/*4^, where *α* is an approximately species-independent proportionality constant. The naming honors Max Kleiber, who first empirically observed this allometric scaling law in the 1930s, and Geoffrey West, who later provided a theoretical explanation for it in 1997 [2]. The relationship is well supported empirically across a wide range of body masses and species, including mammals, birds, reptiles, fish, and invertebrates [11].

The metabolic output rate *B* represents the energy budget available during ontogeny, and part of it can be allocated to support growth — specifically, the generation of new tissues. However, some metabolic energy is also required to sustain and maintain the existing tissue mass at any given time. West’s theory of ontogenetic growth results from applying the law of energy conservation (the first law of thermodynamics), expressed in the form of a differential equation – the equation of motion governing the growth of an organism:

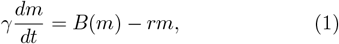

where *γ* and *r* denote the energy costs per unit mass of tissues of growth and somatic maintenance, respectively [4].

The energy cost of maintenance *r* is dominated by the energetic costs of cellular and sub-cellular turnover, which involves the removal and replacement of organelles, cells, and tissues [12]. Such turnover is a routine function in many tissues, including skin cells and mitochondria in metabolically active tissues, serving to remove damaged or exhausted components. Moreover, tissue turnover and regrowth occurs in response to stresses such as injury and infection, where tissue damage is exacerbated by associated inflammatory responses. Indeed, organismal growth is often delayed or inhibited following severe injury or infection, underscoring the significant energetic costs associated with these stress responses[13][14].

In the beginning of the ontogenic growth trajectory, the body mass *m* is small, and most metabolic energy is available to fuel growth (*B*(*m*) ≳ *rm*). However, as body mass increases, the demands for somatic maintenance also rise. The body mass asymptotically approaches a steady state value, *m*_*ad*_, at which point all growth ceases because all available energy is required to maintain the existing tissue mass. Solving Eq. 1 for *m*_*ad*_ yields: *m*_*ad*_ = (*α/r*)^4^. Development time (*t*_dev_) in the model is finite, denoting the time point when development is complete but the natal mass is still well below *m*_*ad*_, the maximal mass of the fully grown animal. In typical cases, it can be shown that: *t*_dev_ ≈4*γ/r* (see the original work [4] for more details and derivations).

### B. The second law and entropic damage

There is no aging in West’s model since the adult state is also the steady state. However, maintaining this steady state necessitates significant metabolic energy being devoted to somatic maintenance - particularly to the ongoing turnover of tissues and organelles. The second law states that not all metabolic energy can be converted into useful work such as reversible pathway activations required for growth and maintenance of tissues; instead, some of this energy will inevitably be lost as heat. While much of this heat is dissipated as waste, a portion will remain within the system in the form of damage or errors, contributing to irreversible damage that accumulates with age. This is simply another way of stating the fundamental reality that no biosynthetic, maintenance, or repair mechanisms operate with perfect fidelity.

In high-dimensional data collected from animals of different ages, the damage manifests as a cluster of features with a linearly increasing mean and variance - the hallmarks of a Poisson stochastic process [7]. Furthermore, the entropic nature of the damage is further corroborated by single-cell DNAm data from aging animals [8]. Although the mathematical properties of stochastic damage are clearly defined, the specific molecular nature of the damage considered here is not critical - the same analysis may be applied to different types of damage, including traditional macromolecular damage, somatic mutations, or loss of hyper-or hypomethylated states in DNA.

The damage accumulation rate (DAR), *R*_*D*_, is a species-specific quantity that quantifies the rate at which molecular and informational damage is generated per cell or per unit of body mass. Thermodynamic considerations dictate that the DAR should be proportional to the mass-specific metabolic activity: *R*_*D*_ = *cB*(*m*_ad_)*/m*_ad_, where *c* is the proportionality coefficient – a species-specific parameter quantifying the number of damaging events that occur on the cellular level per unit of time and metabolic output, also referred to as the thermodynamic fidelity.

Combining the mass and energy properties of the adult organism set by the first law of thermodynamics (see Section II A), we find the following equivalent representations for the DAR:

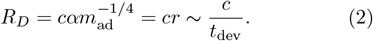

In other words, the cellular DAR is inversely proportional to the body mass to the power 1*/*4. On the other hand, the DAR is also proportional to *r*, the energy cost of maintenance per unit mass of tissue. This relationship implies that for a fixed value of the thermodynamic fidelity factor *c*, more rapid damage accumulation results from faster tissue or organelle turnover (larger r). This is true, for example, in the case of mitochondria where organelle turnover contributes to the generation and expansion of detrimental mtDNA mutations [15][16].

Our next goal is to connect these macroscopic (thermodynamic) arguments to the molecular level (microscopic) properties of aging organisms using the language of non-equilibrium statistical mechanics. To achieve this goal, let us consider the dynamics of an organism’s state as a stochastic process involving transitions among a vast number of metastable states, each representing distinct microscopic configurations. Mechanistically, the transitions between these states represent some form of configuration changes such as macromolecular damage or informational corruption (e.g., mutations, changes in epigenetic state). While molecular networks operating in each state perform broadly similar, each transition results in a small, typically detrimental change to their function. Collectively, the system therefore exhibits progressively decreasing functionality throughout the organism’s life history (see Fig. 1).

**FIG. 1.**
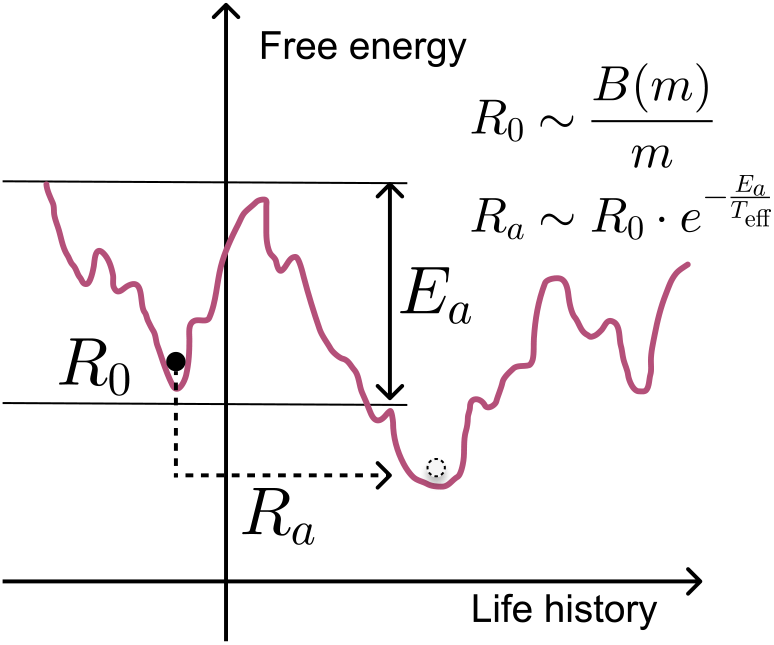
Schematic representation of the configurational transitions in an organism’s life history. The free energy potential surface is shaped by regulatory interactions and is highly rugged. The life history of an organism consists of a sequence of transitions between metastable states, each representing microscopic configurations such as mutations, epimutations, and other forms of state changes of biomolecules. The pathological transitions are relatively rare because they are suppressed by large activation barriers *E*_*a*_. Consequently, the transition rates are low, given by *R*_*a*_ *R*_0_ ∼exp(−*E*_*a*_*/T*), where *R*_0_ is the “attempt” frequency, which is proportional to the basic metabolic rate per cell, and *T* is the temperature -the energy scale available for the transitions.

Assuming that the configurational transition driving the aging process are rare and mostly occur independently, we can focus, at any given time, on a pair of adjacent states representing a configurational transition of immediate interest. For example, this could be the change in the state of a specific methylation site *a* or the transition between the normal and mutated state of a specific nucleotide. These two states are separated by an activation barrier *E*_*a*_.

Since most configurational transitions relevant to aging are slow compared to the time scales of the typical pathway activations driving the modification of macro-molecular states, we conclude that the energy barriers *E*_*a*_ protecting against random transitions are much larger than the energy scale characterizing these fluctuations – the temperature (*T*) (hereinafter we use energy units for temperature variables to avoid excessive repeating factors, such as Boltzmann constant).

In this regime, the transition rates between states are exponentially suppressed: 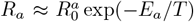,where 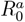 is the “attempt frequency,” the timescale characterizing the frequency of activation transition attempts –a quantity proportional to the metabolic output per cell,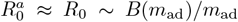. Since the total number of possible configurational transitions is extremely large, the detailed properties of individual transitions and their precise effect are not important. Hence, the average transition rate – the damage accumulation rate, *R*_*D*∼_ *B*(*m*)*c*(*T*)*/m*, can be estimated by integrating over all possible transitions so that the thermodynamic fidelity factor is now temperature-dependent and is given by the integral:

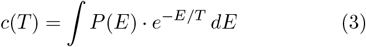

Here, *P* (*E*) represents the statistical properties of the free energy surface defined by the regulatory forces – the effect of the homeodynamic network – characterizing the probability of finding an activation barrier of a given strength *E*.

Eqs. 2 and 3 respectively provide the macroscopic and the microscopic definitions of the thermodynamic fidelity factor *c*. At the macroscopic level, it is the key species-specific constant, determining damage trajectories during aging by relating metabolic output *B*(*m*) to irreversible damage accumulation rate. On a microscopic level, the fidelity factor depends on the structural properties of the free energy potential surface shaped by the regulatory interactions through *P* (*E*) as well as on the temperature *T*. Both properties are subject to evolutionary optimization and therefore may differ between different species. Below, we apply this analytical framework and attempt to identify the relevant parameters from the DNAm data from the Mammalian Methylation Consortium [9] dataset.

### C. Identification of stochastic damage and key pathways by matrix factorization from DNA methylation data

We employed the recently published dataset from the Mammalian Methylation Consortium [9]. We selected samples from 101 evolutionarily distant species with sufficient number of repeats observed in several age-points. We start by estimating pathway activations and the entropic damage burden at different stages of an animal’s adult life directly from DNAm data for each species. To achieve this, we model the observed DNAm patterns as linear combinations of perturbations relative to a reference state. The observed signals in this model originate from biological fluctuations depending on precise physiological status of each animal at the time of sampling, as well as on experimental conditions (including species, sex, age and tissue type).

In the linear regime, the methylation levels 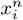 at CpG site *i* in sample *n* can then be expressed as:

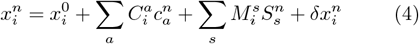

where the vector of covariates 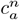 captures information about biological context and experimental conditions, including sex, tissue type, and experimental protocols. The species labels 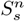 are encoded as one-hot indicator variables, with 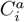 and 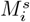 representing the corresponding regression coefficients to be determined from the data. The regression residuals 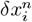 were estimated using linear regression. After fitting the model (4) to account for effects of age, species and tissue, the 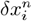 capture additional biological fluctuations, including stochastic pathway activation, unmodeled biological processes, and any remaining batch effects.

To identify co-regulated DNA methylation sites associated with biological pathways (i.e., modes in dynamical systems theory) from the residuals 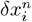,we employed Singular Value Decomposition (SVD), on the residuals 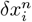:

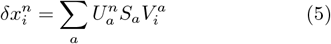

where *a* indexes the different modes, 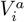 are the loading vectors that characterize the contribution of feature *i* (ith methylation site) to the dynamical mode *a*, and 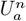 are the scores for each sample *n* and mode *a*. The pathway activations 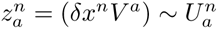 serve as a mode variables or reaction coordinates associated with mode *a* in each sample *n*.

To estimate the signal-to-noise ratio in the dataset, we plotted the distribution of the eigenvalues of the data covariance matrix, given by 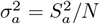,where *N* is the total number of samples in the analysis (see Fig. 2).

**FIG. 2.**
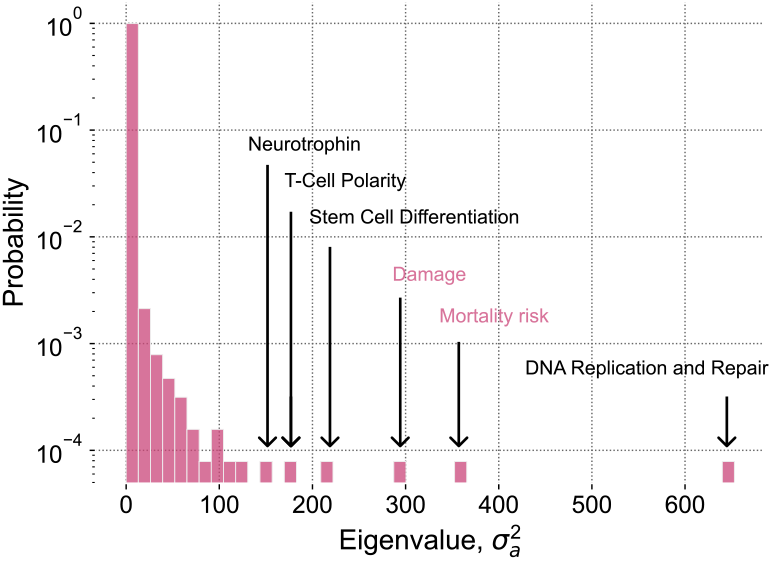
Distribution of normalized singular values from the biological noise covariance matrix. The singular values repre-sent distinct modes of variation in the dataset, each associated with specific biological processes such as DNA replication and repair, stem cell differentiation, T-cell polarity, neurotrophin signaling, and others. Separately, we highlighted the agedependent features: the entropic damage (PC3) and the mortality risk (PC2).

To evaluate the significance of these eigenvalues, we then compared this distribution to that of a random (shuffled) matrix of the same size. The eigenvalues of the shuffled covariance matrix fall within the range of 2.72 · 10^−11^ to 1.66, which aligns with expectations from random matrix theory, where the maximum eigenvalue 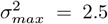 is predicted by Tracy-Widom Law. This comparison demonstrates that 250 eigenvalues exceeding 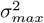 are unlikely to result from random noise, indicating the presence of meaningful biological signals.

The loading vectors of these modes reflect the involvement of individual CpG sites in specific biological processes. To identify the processes involved in these modes we performed functional enrichment study, accounting for the mammalian array background, of genes regulated by CpG sites corresponding to the leading components of the first ten principal components (PCs). The loading vectors for the leading PCs suggest that PC scores are associated with distinct biological processes, such as DNA replication and repair, stem cell differentiation, T-cell polarity, neurotrophin signaling, and others (see Fig. 2).

### D. Characterization of age-dependent processes

Next, we evaluated which of the mode functions (the principle component (PC) scores representing pathway activations (PA) within the proposed model) correlate with the age of the animals from which the samples were taken. To facilitate cross-species comparison, we determined the correlation of PAs with the dimensionless age obtained by rescaling each animal’s age by its species’ maximum lifespan. This analysis revealed that the two principle components (the second and the third) were best correlated to the scaled age (Pearson’s *r* = 0.42, *p* = 2.4· 10^−3^ and *r* = 0.77, *p* = 8.2 · 10^−11^, for PC2 and PC3, respectively), as shown in Fig. 3).

**FIG. 3.**
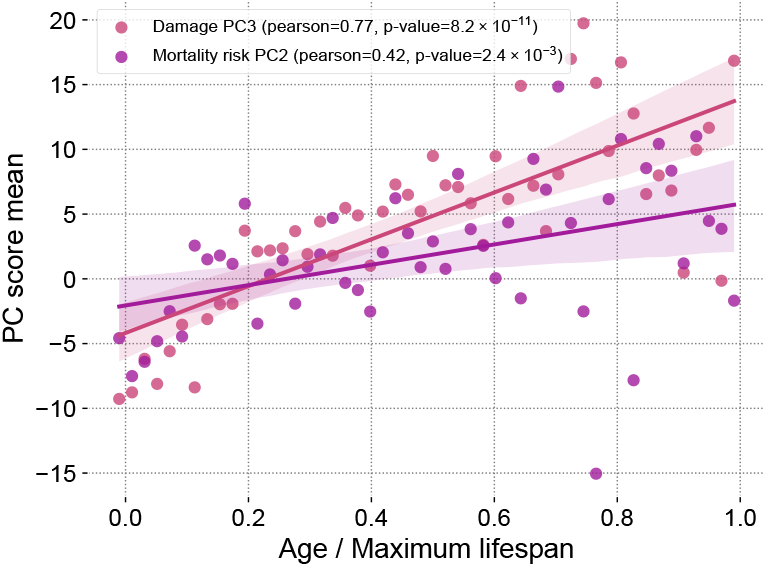
Age dependence of the two leading Principal Component scores (PC2 and PC3) on the (the rescaled) chronological age of the animals.

Importantly, not only does the mean of PC3 increases linearly with age, but also the variance does as well (see Fig. 4). We formally confirmed this observation by determining the correlation between the mean and the variance for PC3, confirming that the two quantities are significantly correlated (*r* = 0.53, *p* = 7.9 · 10^−5^). A stochastic variable displaying a linear increase in mean and a proportionality between its mean and variance is the classic hallmark of data originating from a realization of a Poisson stochastic process – a signature of a variable tracking stochastic damage (see [7, 8] for more examples in other datasets).

**FIG. 4.**
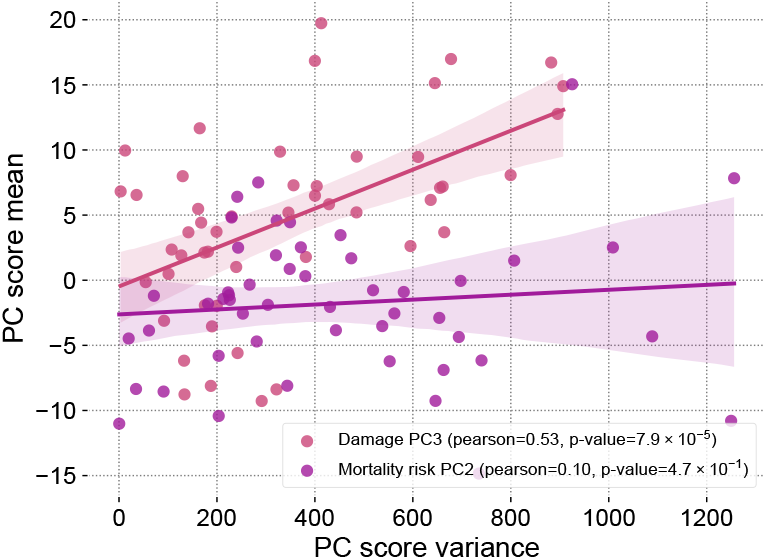
The variance to mean correlation for the leading age-associated PC scores. PC3 has variance proportional to mean -a hallmark of Poisson stochastic process.

We therefore identify the linear in PC3 score in our dataset as metric for the age-dependent cumulative effect of stochastic damage. In this view, PC3 tracks individually rare, stochastic transitions of CpG sites that occur over time as consequence of imperfect maintenance of methylation states. Notably, the Poissonian character of PC3 is unique, with most leading PCs not showing strong age-dependence and PC2 displaying no significant correlation between its variance with its mean (*r* = 0.10 and *p* = 0.47 as shown in Fig. 4).

Having identified PC3 as the signature of stochastic damage accumulation, we investigated the PC3 loading vector components, which are proportional to configuration transition rates, *R*_*i*_, characterizing the dynamics of the methylation transitions at each CpG site *i*. A log-plot of the probability distribution of the vector components of CpG sites in PC3 reveals an exponential distribution of the form: 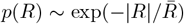 with 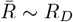 being a constant (see Fig. 5).

**FIG. 5.**
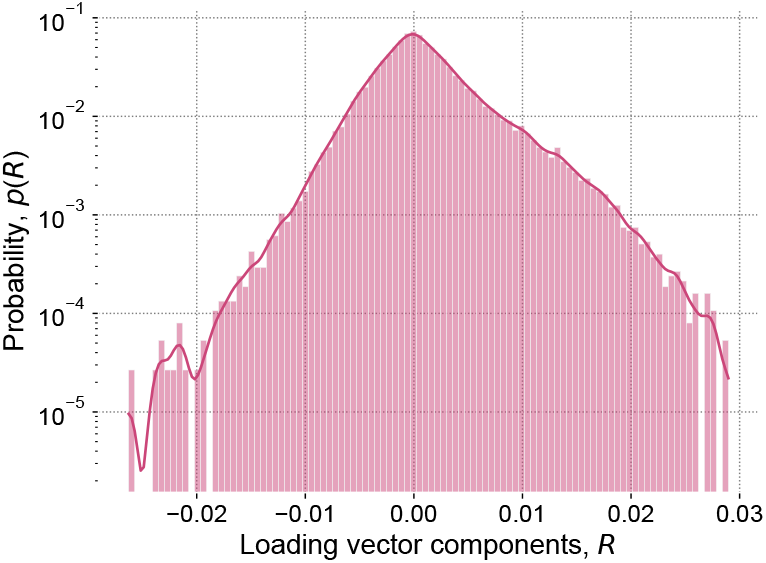
The distribution of the age- and damage-associated PC3 loading vector components - closely related to the distribution of the configuration transition rates *R* (see the explanations in the text).

At the microscopic level, the transition rates are limited by the activation energy barriers protecting individual sites (see Eq. 3 and the relevant description of the microscopic model in Section II B). Since *R*(*E*) = *R*_0_ exp(−*E/T*), the probability distribution of energy barriers protecting individual CpG sites subject to stochastic damage during aging, *P* (*E*), can be obtained by comparing the distributions *p*(*R*)*dR* = *P* (*E*)*dE* leading to:

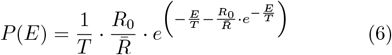

The resulting distribution for the energy barriers *P* (*E*) has the form of a type-I extreme value distribution known as “Gumbel distribution” with location parameter 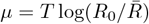 and the scale parameter, both explicitely proporotional to the temperature, *T*. The identification of a Gumbel distribution for activation barriers reveals that the kinetics of the damage accumulation is defined by activation across barriers determined by extreme values across highly rugged and disordered energy landscape.

The variance of PC2, the next leading PC score associated with the rescaled age in the dataset, increases faster than linearly with chronological age. In Fig. 6, we plotted the inverse variance of the leading age-dependent PC scores and observed that the inverse variance of PC2 decreases linearly and approaches zero at the age corresponding to the maximum lifespan. This is a signature of the loss of resilience at the maximum lifespan (see [5, 7] for more examples of this phenomenon in blood markers and DNAm in humans). Thus, we expect that the PC2 score is associated with the risk of death and henceforth refer to PC2 as “Mortality Risk PC2”.

**FIG. 6.**
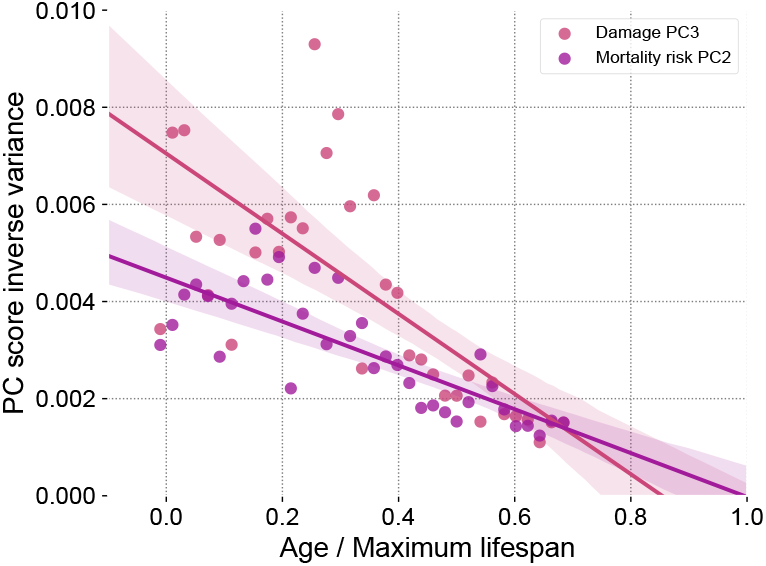
The variance of the leading age-associated PC scores increases with (the rescaled) chronological age of the animals. The inverse variance of PC2 score reveals the loss of resilience (dynamics stability) at the end of life (close to the maximum lifespan).

We conclude this section by estimating the species-specific damage accumulation rate (DAR) as the linear regression slope characterizing the increase of PC3 over time. According to Eq. 2, the DAR should be inversely proportional to the development time, with the proportionality coefficient related to the thermodynamic fidelity factor. We tested this prediction directly by plotting the species-specific *R*_*D*_ against the development time (*t*_*D*_) for each species, finding that the predicted relationship indeed holds true (see Fig. 7). The regression slope on the log-log plot is close to negative one, which aligns with the prediction DAR ∝1*/t*_*D*_. This allows us to establish the logarithm of the “fidelity” factor *c* for each species as the regression residual.

**FIG. 7.**
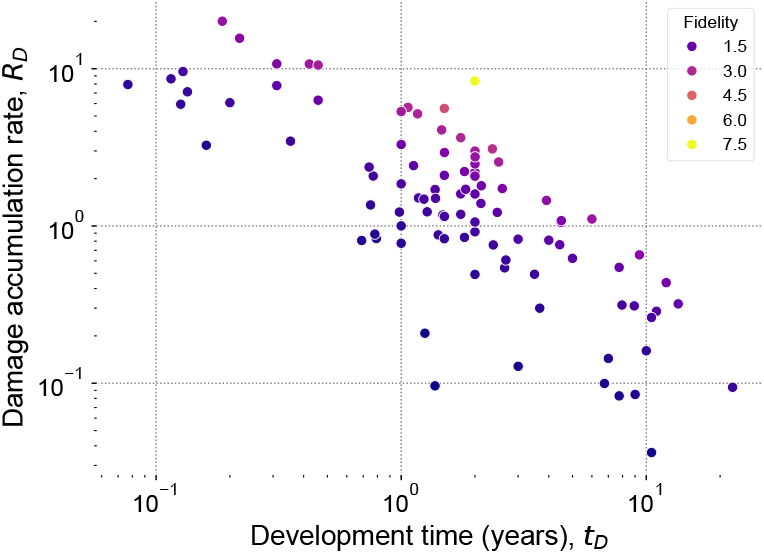
The inverse dependence of the damage accumulation rate (DAR) and the development time.

### E. Universality of pathway activation fluctuations - effective temperature

The availability of multiple measurements from different specimens and various tissues of the same species at different time points enables the investigation of the statistical properties of biological fluctuations in CpG states. Specifically, we focus here on reversible pathway activations – that is, modes other than the stochastic damage signature captured by PC3 – to explore the relationship between fluctuations and dissipation across pathways. We thereby define an effective temperature for each species – a universal (pathway independent) specieslevel parameter controlling the power of physiologically relevant noise.

We assume that pathway activations, represented by the leading principal component (PC) scores 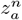 originate from random and reversible biological processes (reversible pathway activations) contributing to the observables 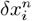. When such fluctuations occur in the presence of homeodynamic regulatory forces, these forces seek to return the system to an equilibrium state. In a linear approximation, the dynamics of the pathway activations follows overdamped Langevin equations:

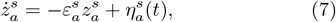

where 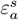 is the species- and pathway-dependent relaxation rates that characterize the rate of restoring of the homeostatic equilibrium in a particular pathway, and 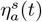 is the stochastic noise (intrinsic or environmental) driving the observed fluctuations. Since pathway activations are presumed to be reversible, this noise satisfies: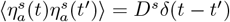, where δ(*t*) is the Dirac delta function [10].

Consequently, the variance of the pathway activation scores can be obtained by solving the Langevin equation (7):

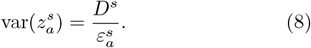

Therefore, we expect that the fluctuations 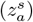 are universally related to dissipation 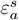 via the power of the noise - a species-specific and pathway-independent proportionality constant playing the role of the effective temperature 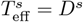.

Living systems are not at equilibrium, and therefore there may be no single temperature factor that characterizes the fluctuation-dissipation relations across all or even most pathways in an organism. In what follows we test whether there is a common factor driving the fluctuations across the slowest pathways with small 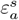 and hence highest variance, according to Eq.(8), which dominate PCA.

The only exception is PC3, which correlates with the total entropic damage and does not represent a specific pathway. Aging occurs on the timescale of the average lifespan of a species, which means it is slower than reversible pathway activations. Reversible pathway activations are subject to evolutionary optimization, requiring them to influence fitness over shorter timescales than the organism’s entire lifespan. Therefore, we assume that aging occurs at timescales much longer than any of the relaxation times 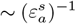,meaning that all pathways have sufficient time to readjust, that is to equilibrate, to the level of slowly increasing damage captured by PC3. Consequently, we do not include PC3 in the analysis aimed at extracting the effective temperature of reversible pathway activation.

The effective temperature and leading relaxation rates can be computed directly if longitudinal data tracking individual subjects through the feature space of interest can be obtained (see, e.g., [5] where we were able to compute *ε*_*a*_ from auto-correlation functions along life histories of the patients and then directly applied Eq. 8 to longitudinal data from blood cell counts and wearable physical activity). Unfortunately, no such data is available as part of the current dataset. However, since, according to Eq. 8:

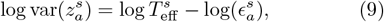

should a common pathway-independent scaling factor (such as effective temperature) driving the pathway activations exist, it could be identified indirectly by a linear matrix factorization of log 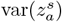.

SVD of log 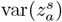 shows that there is indeed a single factor driving the variance of all pathways fluctuations within each species. In fact, the first principal component of the fluctuations explained more than 60% of the variance in PAs, see Fig. S1). Notably, the corresponding loading vector components are not centered on zero (see Fig. S2), and hence, in the first approximation, all reversible pathways increase and decrease their variance together, as expected from Eqs. 8 and 9 (see Fig. 8).

**FIG. 8.**
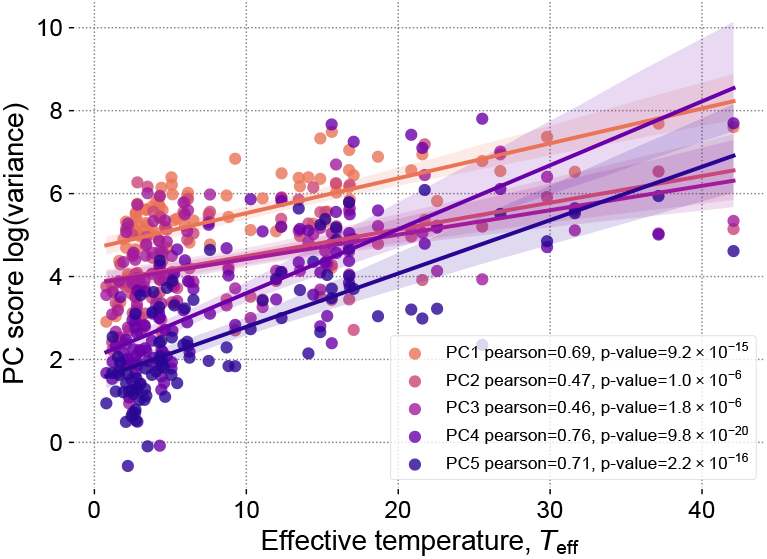
Correlations of the variance of key pathway activations (represented by the variance of the first few PC scores across species) and the effective temperature parameter

Other principal component (PC) scores of pathway activation variances are expected to reflect the variance of the relaxation times of pathways across species, where shorter recovery times reduce the variance of pathway activations).

Therefore, the matrix factorization of the variances of pathway activations in our cross-sectional dataset provides evidence that the power of the biological noise quantified by the the variance of all reversible pathway activations is dominated by a single factor, the first PC of our log-variance analysis. This factor is species-but not pathway-dependent, and can be interpreted as the effective temperature *T*_eff_.

The concept of effective temperature is borrowed from non-equilibrium thermodynamics and may vary on different levels of the organism’s organization. It is, for example, unrelated to the physical body temperature characterizing the fluctuations on the individual molecular level (see Fig. S3 illustrating the lack of correlation between the effective temperature and the body temperature for the same species retrieved from the AnAge database (Pearson’s *r* = 0.05, *p* = 0.68). While physical body temperature determines the reaction rate of chemical reactions, the effective temperature controls the slow dynamics related to pathway activation fluctuations, damage accumulation, incidence of diseases and mortality (see next section).

### F Effective temperature and Gompertz law

Macroscopically, the damage accumulation rate, *R*_*D*_, is proportional to the mass-specific metabolic output and the thermodynamic fidelity, *c*, (see Section A). Neither of these depend on the effective temperature obtained from the analysis of the slowest pathways dominating the variance of physiologically-relevant variables (see Fig. 9). Accordingly, *R*_*D*_ also does not depend on *T*_eff_. In DNAm data, we observed this damage as linear increase in both mean and variance of PC3. While damage therefore increases linearly with age, the risk of death in most animals increases exponentially. This apparent conundrum can be explained by the effects of non-linear coupling leading to the progressive loss of resilience due to the effect of progressive damage accumulation.

**FIG. 9.**
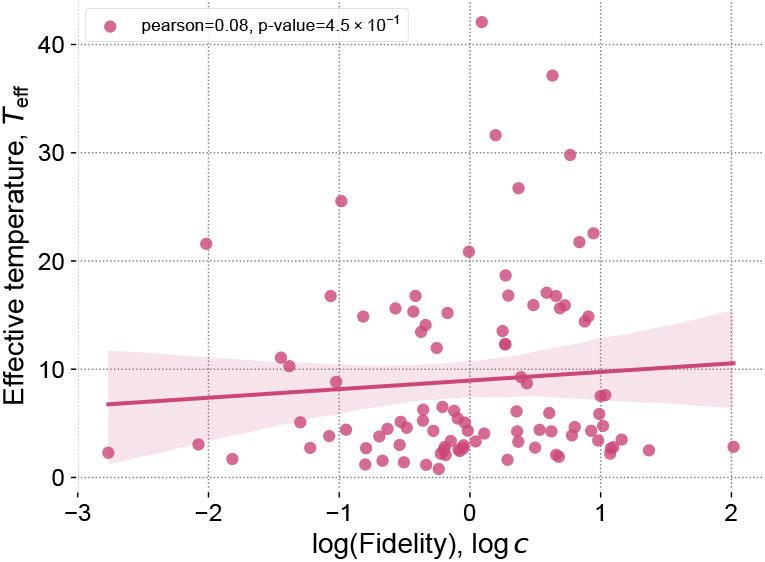
Thermodynamic fidelity, the proportionality coefficient between the damage accumulation rate and the mass-specific metabolic rate does not depend on the effective temperature *T*_eff_

The regulatory networks responsible for maintaining homeostatic functions have evolved to preserve resilience in the face of stochastic damage, resulting in network structures that degrade linearly in performance with over substantioal ranges of damage burden [17]. Microscopically, damage transitions are stochastic, individually rare and each damaging transition results in only a small distortion of regulatory functions. As outlined in [7], the effect a of vast number of uncorrelated events has a cumulative detrimental effect that is independent of the details pertaining to specific transitions with the overall effect being proportional only to the total damage burden, *Z*, which increases, on average, linearly with age, *t*: *Z*(*t*) = *R*_*D*_*t*.

The effect of this damage is to slowly alter the recovery rates *ε*_*a*_, an effect is the most visible for the lowest-frequency (slowest) modes dominating the reversible pathway activations within the PCA. In [5, 7] we observed that in human blood biomarkers and blood DNAm data, there is a mode with the recovery rate approaching zero at the age of 120, roughly corresponding to the maximum lifespan in humans. In a cross-sectional dataset such loss of resilience manifests itself in hyperbolic divergence of fluctuations of pathways strongly associated with age and other than the damage (see Eq. 8). Applied to the current DNAm dataset, we observe a similar pattern for PC2, which inverse variance approaches zero (and hence the variance diverges) close to the maximum lifespan for each species.

In the presence of fluctuations (that is, for any non zero value of *T*_eff_) the stability of the organism state is lost well before the age corresponding to this total loss of resilience. The disintegration of the organism (death) occurs once the basin of homeodynamic stability, formed by the regulatory interactions along the direction coinciding with PC2, becomes shallower than the scale of typical fluctuations as defined by *T*_eff_. In other words, the age-dependent exponential acceleration of mortality occurs due to the linear degradation the protective barriers by the cumulative effect of damage: as described in [7], the activation energy, *E*_0_(*Z*), associated with the least stable mode (here, the mode associated with PC2) in linear approximation is:

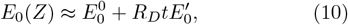

where 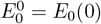, the activation energy in young animals, and 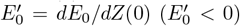 are the first and second order coefficient of Taylor series expansion of the activation barrier strength *E*_0_(*Z*). The mortality in the model can be calculated as the activation transition rate:

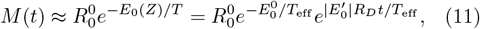

with 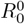 being the pathway specific attempt frequency. Comparing the expression for *M* (*t*) with Gompertz mortality law, we find that the initial mortality rate (IMR) is 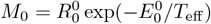,and the associated Gompertz exponent is 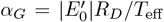 both of which are dependent on the effective temperature due to the fact that the transition rates depend on the magnitude of the fluctuations that are universally scaled by *T*_eff_. Note, that the attempt frequency, 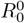, is much larger than the Gompertz exponent, *α*_*G*_.

The maximum lifespan in this model corresponds to the chronological age (or the level of damage) when the protective barrier along with the resilience disappear al-together:

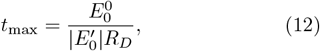

In this analysis, the maximum lifespan is inversely proportional to the damage accumulation rate *R*_*D*_, but, since *R*_*D*_ does not depend on the effective temperature, *T*_eff_, the maximum lifespan is independent of *T*_eff_. By contrast, the average lifespan under Gompertz law is 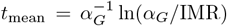 and this can be expressed in terms of *T*_eff_ and *R*_*D*_ as:

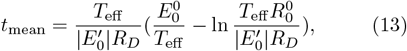

Like the maximum lifespan, the average lifespan is there-for inversely proportional to the damage accumulation rate, but, unlike the maximum lifespan, the average lifespan is explicitly dependent on the effective temperature. Given that the quantity under the log is the large ratio 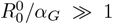,the second term is always positive, and hence the average lifespan is always smaller than the maximum lifespan, with the difference between the two depending on the effective temperature:

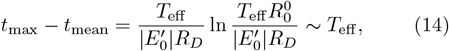

Therefore, the effective temperature *T*_eff_ controls the shape of the survival curve with diminishing temperature corresponding to increasing “squaring off” the curve. The survival curve becomes perfectly square, with mean and the maximum lifespan coinciding exactly, *t*_mean_ = *t*_max_, in the theoretical limit of vanishing fluctuations, that is when: *T*_eff_ = 0.

To demonstrate that the effective temperature obtained from DNAm data indeed controls the difference between the mean and the maximum lifespan in different mammalian species, we next plotted the ratio of the maximum to the average lifespan using lifetables data from [18] against the temperature estimates from our work (see Fig. 10). As predicted, the ratio of the lifespans is positively associated with the effective temperature (Pearson’s *r* = 0.36, *p* = 0.05).

**FIG. 10.**
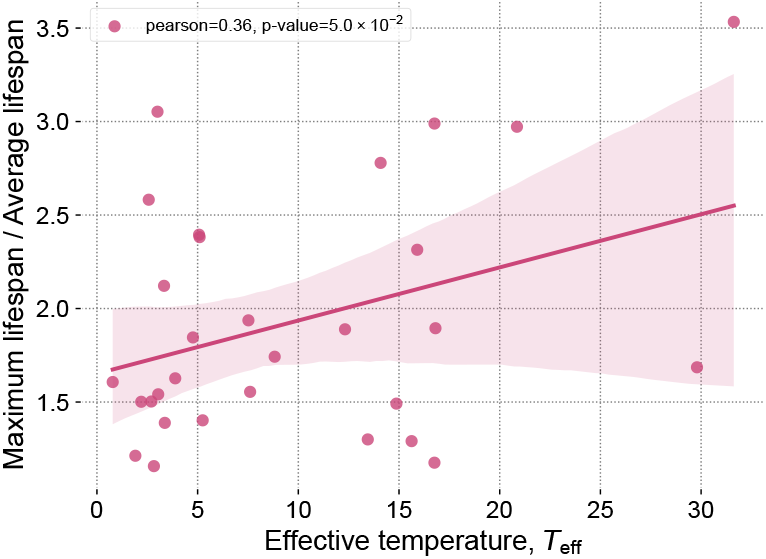
The effective temperature *T*_eff_ controls the rate of actuarial aging (the senescence rate) obtained from fits of mammalian lifetables from [18].

The theoretical description of the aging process proposed here also offers a novel link to the earlier phenomenological Strehler-Mildvan theory of aging [19]. The progressively decreasing protective barrier strength *U* (*Z*) here plays the role of the phenomenological ‘vitality’ parameter introduced by Strehler and Mildvan. However, unlike this earlier concept, the microscopic energy barriers are in principle observable, opening up novel avenues for intervention. Moreover, according to Eq. 11, we expect that:

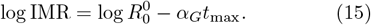

This is nothing but the well-known Strehler-Mildvan correlation, first discussed over 60 years ago in the same earlier work. Our model provides a microscopic explanation for this correlation and suggests that it must hold if the structural properties, such as 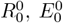 and 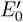 are fixed by slowly evolving biology while the temperature *T*_eff_ changes due to faster acting (e.g. social or technological) causes.

## III. DISCUSSION

In this Letter, we investigate the implications of systematically applying the first (energy conservation) and second (entropy increase) laws of thermodynamics to the processes of development and aging. It has long been recognized that the combination of the Kleiber-West law, which dictates the scaling of basic metabolic output with body mass [2], and energy conservation leads to universality of ontogenic growth trajectories across species. Additionally, these principles impose constraints on development time and adult body mass as functions of energy allocation toward maintenance, tissue repair and replacement [4].

However, the existing model developed by West does not account for aging, as the fully grown organism is considered a dynamically stable equilibrium state. Building on our previous work [5, 7], we propose that the long term stability of the adult state is limited by the cumulative effect of progressively increasing micro-insults, that is, the accumulation of irreversible damage. We therefore propose to extend West’s model by incorporating aspects of the second law of thermodynamics and applying kinetic arguments to introduce a novel analytical framework to analyze the role of species-specific damage accumulation rate, *R*_*D*_, in aging. The resulting framework provides testable hypotheses, including a formalism to quantitatively determine the effect of *R*_*D*_ on the maximum and the average lifespan of species.

At the macroscopic, that is, at the level of organism, we assume that the damage accumulation is proportional to the basal metabolic output per cell. The proportionality coefficient, the thermodynamic fidelity factor, controls the proportion of available energy (given by the basal metabolic rate) dissipated throughout an organism’s life that is transformed into irreversible damage. The relationship between mass-specific basal metabolic rate and longevity is, of course, very well established (see, e.g., the most recent [20], which demonstrates causal connections between basal metabolic rate and epigenetic age acceleration through bidirectional Mendelian randomization and mediation analysis). The framework presented here is different from the rate of living or oxidative stress theories of aging, in that damage is conceptualized as a fundamental consequence of unattainability of perfect conversion of heat into work (second law of thermodynamics), rather than as a direct result of specific forms of macromolecular, e.g. oxidative damage. Damage here refers to any consequence of thermodynamic inefficiency that results in residual energy in a high-entropy state. This generalized damage can manifest as corruption of information (mutations, change in DNAm state), macro-molecular (e.g., oxidative, adducts) or at systems level (e.g., changes in physiological set points).

According to G. West’s argument [4], in a fully grown organism all metabolic output is used for organismal and tissue maintenance and hence the metabolic output per unit of mass equals the energy expenditure required for maintenance per mass. Therefore, the damage accumulation rate is proportional to the same energetic cost of tissue turnover and repair, primarily driven by cellular and sub-cellular renewal (turnover). The proportionality constant, the thermodynamic fidelity factor, can therefore be interpreted as a species-specific probability that any given turnover or repair event will produce irreversible (entropic) damage. This interpretation underscores the idea that, while repair mechanisms are essential for maintenance and survival, especially given the constant physical assault that characterizes the natural environment, the same process also carry an inherent risk of contributing to entropic aging.

Like all biological traits, the rapidity and fidelity of biosynthetic and maintenance pathways is shaped by evolutionary pressures. For instance, the accuracy of DNA polymerases directly influences the rate at which somatic mutations accumulate, with significant implications for cellular senescence and cancer [21]. Studies on mutator mice, specifically the mtDNA mutator mouse model characterized by a proofreading deficiency in mitochondrial DNA polymerase (POLG), have shown that these mice accumulate high levels of point mutations, leading to premature aging phenotypes such as weight loss, reduced fertility, and decreased lifespan [22, 23]. However, higher fidelity in these processes typically comes at the cost of substantial energetic and performance detriments. Proofreading polymerases, for example, while more accurate, require additional energy and exhibit lower processivity compared to their less accurate counterparts; this is exemplified by the differences between Taq polymerase and Deep Vent polymerase [24].

Balancing repair and recovery capacity and energetic cost of these processes with the entropic damage accumulation rate is a classic example of an evolutionary trade-off between short term survival in a hostile environment and long term stability (entropic aging). In light of Haldane’s and Medawar’s concepts of the “selection shadow,” it follows that evolutionary optimization will result in species-specific trade-offs between cell turnover, molecular fidelity, resource use, and performance [25]. As a result, we can expect entropic damage to accumulate as a consequence of finite fidelity of maintenance in most adult animals, albeit at rates that vary across species. Interestingly, species living in hostile environments—characterized by high turnover and likely short lifespans—may still provide instructive examples of ways to achieve exceptional thermodynamic fidelity.

Next, we strengthen the thermodynamic argument, extending our analysis to the microscopic level by proposing a semiquantitative model of the fidelity factor. This extension is based on the theoretical model from [7]: the damage accumulation rate is determined as the average configurational transition rates, characterizing rare events such as mutations, epimutations, or transitions into altered (pathological) states, separated from the normal operational states by activation barriers of various heights. The damage accumulation rate in this picture is an average quantity that depends separately on the structural and thermodynamic properties of the organism: the statistical properties of the free energy surface, shaped by the regulatory interactions (homeodynamic network) and by an effective temperature of the organism.

To bridge the macroscopic (thermodynamic) and microscopic (molecular) perspectives, we leveraged the newly published longitudinal DNA methylation (DNAm) data from 348 mammalian species separated by large phylogenetic distances[9]. We performed a simple normal mode analysis using the SVD matrix factorization technique on these data, identifying modules of correlated DNAm features. While most of these signals (modes) originate from reversible pathway activation of key biological processes (organism-wide pathways).

The ability of PCA or related clustering algorithms to identify physiologically relevant modules should not be surprising since PCA can approximate the inverse Hessian matrix when a single energy well is sampled, which is closely related to normal mode analysis. The examples span multiple areas of applications ranging from molecular dynamics (see, e.g., [26]) to identification of modules clustering stocks according to their respective industries in financial market data [27]. In [9], unbiased clustering of individual cytosines in the dataset used in this work led to identification of 31 modules related to primary traits including age, species lifespan, sex, adult species weight, tissue type and phylogenetic order.

The leading age-dependent PC score (PC3) has a property of a Poissonian random variable - having both its mean and variance increase linearly with chronological age. Intriguingly, for different species, the rate of this increase is inversely proportional to the maximum lifespan of the species from which the data was derived. Following [7, 8], we interpret the Poissonian PC3 score as the signature of irreversible damage accumulation - the measure of entropic damage. We then use the “linear” PC3 score to estimate the damage accumulation rate, *R*_*D*_, for each species. We next confirmed the expected inverse association of *R*_*D*_ with the development time (see Eq. 2 of the macroscopic model) and used the regression residual of this fit to estimate the thermodynamic fidelity factor for each species.

The loading vector components of the principal component (PC3) associated with linear (entropic) damage represent configurational transition rates for CpG sites that collectively define damage accumulation. The distribution of these loading vector components is non-Gaussian, exhibiting heavy exponential tails, meaning that the energy barriers protecting different CpG sites is distributed non-Gaussian. Further analysis of this microscopic model showed that this exponential distribution of configuration transition rates corresponds to a Gumbel distribution in the energy of activation barriers—a type of universal distribution commonly encountered in extreme value statistics (e.g., the distribution of maximum or minimum values in a series of random variables, see [10]).

Finding that the distribution of activation barriers in relation to kinetic of damage accumulation follows a Gumbel distribution provides intriguing insights into the underlying dynamics and structural characteristics of regulatory networks relevant to aging. In our context, this may suggest a rugged and highly heterogeneous energy landscape with a wide range of energy barriers separating them. The applicability of the Gumbel distribution often relies on the assumption that the activation barriers are independent or only weakly correlated. The emergence of the Gumbel distribution from the CpG data may imply that the barriers are not systematically related to each other, allowing the extremes to dominate the distribution. In systems where activation barriers follow a Gumbel distribution, the kinetics are often controlled by the highest barriers. This means that the rate-limiting steps along the aging trajectory are rare, high-energy transitions – most probably unlikely, simultaneous failures in highly redundant systems. Consequently, aging emerges as a thermodynamically irreversible phenomenon, as reversing these unique combinations of random events would demand an impractical degree of control at the microscopic level.

The availability of multiple replicates from the same species allowed us to next analyze the statistical properties of the biological noise associated with reversible pathway activations and its differences between species. We identified a common factor that explains most of the variance in biological fluctuations across the leading (most variable) pathways. The presence of such a factor suggests that the variability in pathway activation is not merely an inherent property of specific pathways but reflects the redistribution of energy between the collective degrees of freedom (pathways) due to strong interactions between the constituents of biological systems. The common scaling factor in biological fluctuations emerges as a global (species-specific) thermodynamic property of mammalian regulatory networks - a property that is best described as the effective temperature of the species.

The connection between effective temperature and pathway activation variance is analogous to the Einstein relation between fluctuations and dissipation in systems close to equilibrium [10], an observation that highlights the broad applicability of core thermodynamic principles to biological systems. While in equilibrium, the temperature parameter is truly global, aging organisms are clearly in a non-equilibrium state and therefore the existence of a global temperature parameter may be seen as surprising. Certainly, at the scale corresponding to the kinetics of individual chemical reactions, the relevant temperature parameter should be the ambient or body temperature. However, the effective temperature defined above pertains to another scale (fluctuations in reversible pathway activation) and should not be confused with the physical body temperature as *T*_eff_ does not correlate with body temperature.

The concept of the effective temperature helps explain how the linear damage accumulation translates into the exponential age-dependent acceleration of mortality. It defines both the initial mortality rate and the Gompertz exponent, which are the key demographic (or actuarial) parameters characterizing the aging process. In [7] we proposed that the age-dependent loss of the resilience occurs due to the compounding effects of linearly acquired damage. The end of life in this picture comes when the stability of the equilibrium state is lost upon reaching the limiting age- the maximum lifespan, where the protective barrier is lost and the recovery into the functional state is no longer possible [5]. In the current work, we observed several PCs (modes) representing features other than the damage that also dependent on age. The variance of the leading of these aging signature, PC2, diverges at about the maximum lifespan for each species. This observation suggests that at this age, the progressive loss of resilience results in a vanishing recovery rate along PC2. In other words, the homeostatic equilibrium, defined by the basin of attraction along PC2 is completely lost at the maximum lifespan. This phenomenon has been previously observed for humans in longitudinal datasets of clinical blood markers [5] and DNAm from blood samples [7]. The current work suggest this may be a universal feature of aging in long lived mammals.

Death will occur with certainty when all resilience along any physiological pathway is completely lost. Alternatively, and in fact far more likely, stability may also be lost in the presence of fluctuations able to drive activation over a still present (but insufficient) protective barrier. The power of noise - quantified by the effective temperature - thus determines the age of such premature loss of stability and hence the difference between the maximum and the average lifespan. While neither the rate of aging nor the maximum lifespan are temperature dependent, the mortality trajectory (and, by extension, disease risks) in fact exhibit strong dependence on the effective temperature.

The effective temperature, is a universal properties of the slowest pathways dominating the variance of biological signals and represents the biological processes responsible for the hallmarks of aging and diseases. The presence of a Gumbel distribution, easily explainable by thermal activation transitions, suggests the existence of a distinct effective temperature operating on the rate scales relevant to damage accumulation, which is different from the previously defined effective temperature *T*_eff_. This indicates that the manifestations of aging and diseases, on one hand, and irreversible entropic aging (damage accumulation), on the other, are driven by distinct physical and biological processes on different levels of the organism organization.

The multiscale theoretical framework proposed in this work supports the earlier and purely phenomenological theory of aging by Strehler and Mildvan, formulated in analogy to chemical reaction kinetics theory in [19]. Within the theory, every organism is characterized by a linearly decreasing vitality (physiological potential) and exponential acceleration of mortality is exponentially increasing probability of death due to activation transitions against vitality-dependent protective barriers at finite temperature. Hence, our model provides microscopic definitions of the key parameters of Strehler-Mildvan theory: the PC2 score is the reaction coordinate characterizing the decay of the metastable state; vitality is the activation energy of the smallest protective barrier progressively affected by damage; and the effective temperature, a property of fluctuations of key biological pathways, controls the probability of activation transitions.

Somewhat counterintuitively, a reduction in the effective temperature leads to accelerated actuarial aging (shorter mortality rate doubling time). This phenomenon has in fact been observed in human populations over the past few centuries, where the correlation between the initial mortality rate (corrected for age-independent mortality) and the actuarial aging rate aligns with the Strehler-Mildvan correlation and corresponds to the “squaring off” or “rectangulation” of the survival curve (see, e.g., [28]). In our model, this correlation and hence the historical trends in demographic parameters, corresponds to a line generated by decreasing effective temperature under a constant rate of aging, and therefore a fixed maximum lifespan. The model therefore explains Strehler-Mildvan correlation by postulating that human interventions upon their living conditions and lifestyle have effectively lowered their own effective temperature, suggesting that this parameter can be modulated by lifestyle, environmental and medical means.

Thus, the effective temperature is a new independent control variable that does not influence the rate of aging itself (and consequently does not alter the maximum lifespan). It fundamentally differs from aging clocks by capturing shared properties of biological fluctuations rather than changes in levels (location) of particular age- and health-related markers (see, e.g.,[29] We also note that the role of the effective temperature in regulating the average lifespan without affecting the maximum lifespan, is not the same as the association between increasing noise parameters with biological age or with the rate of aging - noted and characterized in [7, 31, 32]. Moreover, the latter circumstance may not be easily made actionable, since it is merely a property of the entropic damage being a Poissonian random variable, characterized (by definition), by a strong correlation between its mean and its variance, as demonstrated in [7] and in this work.

The effective temperature plays a crucial role in determining healthspan by reshaping the survival curve. As effective temperature decreases, the average lifespan increases, approaching the maximum lifespan for the species. This implies that interventions targeting the effective temperature would broadly impact the risks associated with chronic diseases and mortality across pathways and systems, precisely because the effective temperature is a shared scaling parameter across key physiologically relevant pathways. We therefore advocate for the development of new medical interventions aimed at reducing effective temperature as a promising biotechnology strategy to alleviate the burden of chronic diseases.

It should be emphasized, however, that reducing effective temperature does not impact the underlying rate of aging. Consequently, efforts to extend healthspan through effective temperature modulation will not inherently slow the aging process and hence would not intercept most of the functional decline associated with aging. Aging in mammals appears to be a thermodynamically irreversible process relying on the most fundamental biological mechanisms. Further progress requires the investigation of biological mechanisms behind thermodynamic fidelity that could potentially be targeted pharmacologically. This should open avenues for interventions aimed at modulating the underlying drivers of mammalian aging as a meaningful strategy to slow down the aging process and produce a significant extension of human life.

## IV. ACKNOWLEDGEMENTS

We would like to thank S. Matrenok, N. Tolwinski and D. Igumnov for insightful discussions and comments on the manuscript.

## Appendix S1

**FIG. S1.**
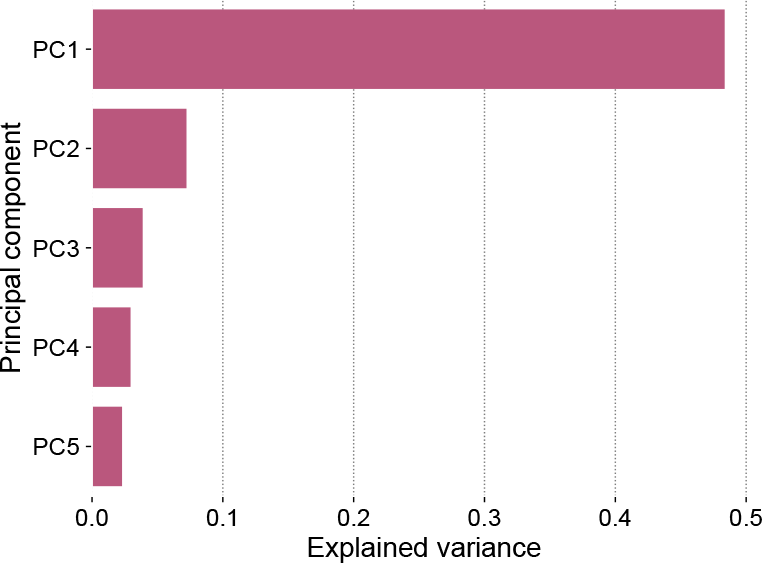
The variance of log-transformed noise estimates, log 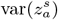,in pathway activations across the species. The figure reveals the dominant factors controlling the power of biological noise.

**FIG. S2.**
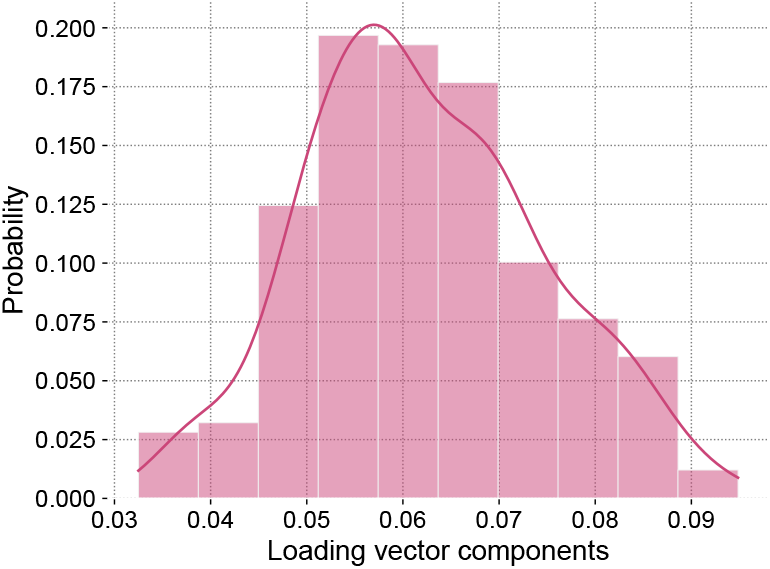
The distribution of the leading loading vector components from the factorization of the log-transformed variability of pathway activations, log 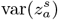.

**FIG. S3.**
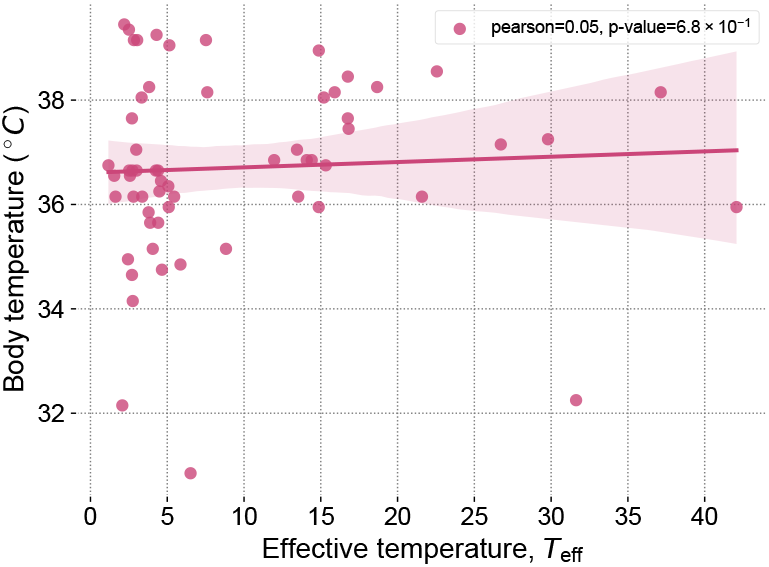
The effective temperature *T*_eff_ does not correlate with the body temperature.

